# Grow with the flow: a latitudinal cline in physiology is associated with more variable precipitation in *Erythranthe cardinalis*

**DOI:** 10.1101/080952

**Authors:** Christopher D. Muir, Amy L. Angert

**Affiliations:** Biodiversity Research Centre and Department of Botany, University of British Columbia, Vancouver, BC, Canada; Department of Zoology, University of British Columbia, Vancouver, BC, Canada

**Keywords:** local adaptation, cline, photosynthesis, growth rate, monkeyflower, *Mimulus*

## Abstract

Local adaptation is commonly observed in nature: organisms perform well in their natal environment, but poorly outside it. Correlations between traits and latitude, or latitudinal clines, are among the most common pieces of evidence for local adaptation, but identifying the traits under selection and the selective agents is challenging. Here, we investigated a latitudinal cline in growth and photosynthesis across 16 populations of the perennial herb *Erythranthe cardinalis* (Phrymaceae). Using machine learning methods, we identify interannual variation in precipitation as a likely selective agent: Southern populations from more variable environments had higher photosynthetic rates and grew faster. We hypothesize that selection may favor a more annualized life history – grow now rather than save for next year – in environments where severe droughts occur more often. Thus our study provides insight into how species may adapt if Mediterranean climates become more variable due to climate change.

## Introduction

Local adaptation has been documented within numerous species; populations generally 2 have higher fitness in their native environment, but perform poorly outside it (Schluter, 3 2000; Leimu and Fischer, 2008; Hereford, 2009). However, the prevalance of local adaptation remains difficult to assess because researchers rarely test for local adaptation unless 5 there are obvious phenotypic or environmental differences (but see Hereford and Winn 6 2008). When local adaptation occurs, it frequently leads to clines in both phenotypes and allele frequencies when selection varies over environmental gradients (Huxley, 1938; Endler, 1977; Barton, 1999). Phenotypic differences between populations along a cline often have a genetic basis and can be studied in a common garden (Turesson, 1922; Clausen et al., 1940; Hiesey et al., 1942). Despite a long history of studying local adaptation and clines, it remains challenging to identify exactly which traits are under selection and which differ for nonadaptive reasons. In particular, the role that physiological differences play in local adaptation is poorly understood, despite the fact that physiology is frequently assumed to explain adaptation to the abiotic environment. A related problem is identifying which of the myriad and often covarying aspects of the environment cause spatially varying selective pressures.

When populations are locally adapted, reaction norms for fitness will cross, such that local genotypes have higher fitness than foreign genotypes and rank orders change across environments (Kawecki and Ebert, 2004). The traits that underlie local adaptation, however, need not mirror this pattern. Populations can have fixed genetic differences conferring trait values that are adaptive at home but neutral or maladaptive away. Alternatively, the ability to plastically respond to a particular environment or the magnitude of response to an environment could be adaptive. We distinguish between these patterns of adaptive trait differences by referring to ‘genetic variation’ in trait means and ‘genetic variation in plasticity’, respectively. Genetic variation in plasticity is synonymous with genotype-byenvironment interactions, or simply (G×E). Genetic variation in trait means and plasticity are both involved in adaptation. For example, genetic variation in photoperiod responses (Blackman et al., 2011) and developmental rate (Stinchcombe et al., 2004) allow organisms to properly time their life history with the local environment. Conversely, sun and shade plants do not have intrinsically higher or lower rates of carbon assimilation, but rather, genetic variation in plasticity cause sun plants to assimilate more under high light and shade plants under low light (Givnish, 1988). In plants especially, we know little about the prevalence and adaptive significance of variation in fundamental physiological traits like photosynthesis and their impact on plant performance (Flood et al., 2011).

A basic approach to identify candidate traits underlying local adaptation is to find associations between traits and environments. Either genetic variation in trait means and/or plasticity should vary clinally along environmental gradients. Indeed, clines in ecologically important traits are widespread in nature (Endler, 1977) and often adaptive, but in most cases the selective agent is unknown. For example, in *Drosophila* numerous latitudinal clines exist for traits like thermal tolerance (Hoffmann et al., 2002), body size (Coyne and Beecham (1987) and references therein), and life history (Schmidt et al., 2005). Some *Drosophila* clines have evolved multiple times (Oakeshott et al. (1982); Huey et al. (2000), see also Bradshaw and Holzapfel (2001)) or shifted in response to climate change (Umina et al., 2005), evincing climatic adaptation. Similarly, plant species exhibit latitudinal clines in traits like flowering time (Stinchcombe et al., 2004), cyanogenesis (Kooyers and Olsen, 2012), leaf morphology (Hopkins et al., 2008; Stock et al., 2014), and drought resistance (Kooyers et al., 2015) that likely relate to climatic variation.

Despite the fact that latitudinal clines have been studied for a long time, latitude *per se* cannot be a selective agent. Latitude may be strongly correlated with one or two key climatic variables, such as temperature, precipitation, or growing degree-days. Latitude may also correlate with the strength of biotic interactions (Schemske et al., 2009) or other nonclimatic aspects of the environment, though as we explain below, we do not yet have compelling data that these are important in our study system. Hence, we focus on whether latitude could be an effective proxy for an underlying climatic driver, in which case we would expect a yet stronger relationship between traits and the key climatic variable(s) driving selection. Alternatively, latitude may be more strongly related to traits than any single climatic variable for at least two reasons. First, latitude may be correlated with several climatic agents of selection that are individually weak, but add up to a strong latitudinal cline. Alternatively, gene flow among neighbouring populations could smooth out local climatic effects, since alleles will experience selection across populations linked by migration (Slatkin, 1978; Paul et al., 2011; Hadfield, 2016). We refer to this as the ‘climatic neighborhood’. For example, in mountainous regions average temperature at a given latitude varies widely, but in aggregate, a lower latitude set of populations will experience warmer climate than a higher latitude one. Thus, any particular low latitude population would be warm-adapted, even if it was located in a cooler (e.g. high elevation) site. Because many climatic factors vary latitudinally, and which climatic factors vary latitudinally changes over the earth’s surface (e.g. coastal vs. continental), dissecting the evolution of latitudinal clines across many species will help identify generalities, such as whether thermal tolerance maxima or seasonal timing is more important (Bradshaw and Holzapfel, 2008), and whether local or regional climate shapes selective pressures.

In this study, we investigated two major questions: 1) whether genetic variation in physiological trait means or plasticity corresponds with latitude; and 2) what climatic factor(s) could plausibly be responsible for latitudinal clines. Within question 2, we tested three hypotheses outlined in the previous paragraph: latitudinal clines are explained by a single dominant climatic factor, multiple climatic factors, or the climatic neighborhood experienced by nearby population connected through gene flow. These hypotheses are not mutually exclusive since, for example, single or multiple factors in a climatic neighborhood may lead to latitudinal clines. We focused on climate because climate often determines where species are found and also can exert strong selection on populations within species. We acknowledge that other abiotic and biotic factors could contribute to selection and the overall pattern of local adaptation. Furthermore, there is a compelling need to know how populations are (or are not) locally adapted to climate so as to predict how they will respond to climate change (Aitken and Whitlock, 2013).

We examined these questions in *Erythranthe cardinalis* (formerly *Mimulus cardinalis* [Ne-som 2014]) because linking physiological traits to potentially complex patterns of local adaptation requires integrating multiple lines of evidence from comparative, experimental, and genomic studies under both lab and field conditions. Many classic and contemporary studies of local adaptation use *Mimulus sensu* lato species because of their natural history, easy propagation, and genetic/genomic resources (Clausen et al., 1940; Hiesey et al., 1971; Bradshaw and Schemske, 2003; Wu et al., 2008; Lowry and Willis, 2010; Wright et al., 2013). Yet, there is a deficiency of links between local adaptation and physiological mechanisms (Angert, 2006; Angert et al., 2008; Wu et al., 2010; Wright et al., 2013). We measured genetic variation in trait means and plasticity in response to temperature and drought among 16 populations distributed over 10.7°of latitude. We found a latitudinal cline of trait means in photosynthesis and growth, but little evidence for variation in plasticity. Interannual variation in precipitation and temperature are associated with this axis of variation, suggesting that climatic variance rather than mean may be an important driver of local adaptation in *E. cardinalis*. The climatic neighborhoods around populations explained trait variation better than local climate, indicating that latitudinal clines may be common because latitude integrates effects of selection on populations connected through gene flow. We place these findings in the context of life history theory and consider future directions in the Discussion.

## Material and Methods

Data and annotated source code to reproduce these analyses and manuscript are available on GitHub (https://github.com/cdmuir/card-cline).

### Population Selection

*E. cardinalis* is a perennial forb native to the Western US (California and Oregon). It is predomintantly outcrossing, self-compatible, and pollinated primarily by hummingbirds. We used 16 populations from throughout the range of *E. cardinalis* (Table 1). These populations were intentionally chosen to span much of the climatic range of the species based on all known occurrences (see below). Seeds were collected in the field from mature, undehisced fruit left open for 2-4 weeks to dry, then stored at room temperature. To control for maternal effects, we grew a large number of field-derived seeds in the greenhouse and generated seed families for this experiment by haphazardly crossing individuals from the same population. We selected seed families to maximize the number of field-derived individuals represented. Thus, we used seeds from 154 greenhouse-derived seed families, 4–12 (mean = 9.6, median = 12) families per population.

**Table 1:** Latitude, longitude, and elevation (mas = meters above seal level) of 16 focal populations used in this study.

| Name | Latitude | Longtiude | Elevation (mas) |
| --- | --- | --- | --- |
| Hauser Creek | 32.657 | -116.532 | 799 |
| Cottonwood Creek | 32.609 | -116.7 | 267 |
| Sweetwater River | 32.9 | -116.585 | 1180 |
| Grade Road Palomar | 33.314 | -116.871 | 1577 |
| Whitewater Canyon | 33.994 | -116.665 | 705 |
| Mill Creek | 34.077 | -116.873 | 2050 |
| West Fork Mojave River | 34.284 | -117.378 | 1120 |
| North Fork Middle Tule River | 36.201 | -118.651 | 1314 |
| Paradise Creek | 36.518 | -118.759 | 926 |
| Redwood Creek | 36.691 | -118.91 | 1727 |
| Wawona | 37.541 | -119.649 | 1224 |
| Rainbow Creek | 37.819 | -120.007 | 876 |
| Middle Yuba River | 39.397 | -121.082 | 455 |
| Little Jamison Creek | 39.743 | -120.704 | 1603 |
| Deep Creek | 41.668 | -123.11 | 707 |
| Rock Creek | 43.374 | -122.957 | 326 |

### Plant propagation

On 14 April, 2014, 3-5 seeds per family were sown directly on sand (Quikrete Play Sand, Georgia, USA) watered to field capacity in RLC4 Ray Leach cone-tainers placed in RL98 98-well trays (Stuewe & Sons, Inc., Oregon, USA). We used pure sand because *E. cardinalis* typically grows in sandy, riparian soils (A. Angert, pers. obs.). Two jumbo-sized cotton balls at the bottom of cone-tainers prevented sand from washing out. Cone-tainers sat in medium-sized flow trays (FLOWTMD, Stuewe & Sons, Inc., Oregon, USA) to continuously bottom-water plants during germination in greenhouses at the University British Columbia campus in Vancouver, Canada (49°15’ N, 123°15’ W). Misters thoroughly wetted the top of the sand every two hours during the day. Most seeds germinated between 1 and 2 weeks, but we allowed 3 weeks before transferring seedlings to growth chambers. We recorded germination daily between one to two weeks after sowing, and every 2-3 days thereafter. On 5 May (21 days after sowing), we transferred seedlings to one of two growth chambers (Model E-15 Conviron, Manitoba, Canada). We thinned seedlings to one plant per conetainer, leaving the center-most plant. 702 of 768 (91.4%) had plants that could be used in the experiment. We allowed one week at constant, non stressful conditions (day: 20°C, night: 16°C) for plants to acclimate to growth chambers before starting treatments. The initial size of seedlings, measured as the length of the first true leaves, did not differ between populations, families, or treatments (Table S1).

**Table S1:** Initial size of seedlings did not vary among Populations, Families, or Treatments. We used a censored Gaussian model of initial size at the outset of the experiment (longest leaf length of the first true leaves). The model was censored because we could not accurately measure leaves less than 0.25 mm with digital callipers (217 of 702, 30.9%, were too small). We fit models using a Bayesian MCMC method implemented using the MCMCglmm function with default priors in the R package **MCMCglmm** version 2.17 (Hadfield, 2010). We estimated the posterior distribution from 1000 samples of an MCMC chain run for 10^5^ steps after a 10^4^ step burn-in. We used step-wise backward elimination procedure to find the best-supported model according to Deviance Information Criterion (DIC).

| Model | Random | DIC |
| --- | --- | --- |
| Population + Water + Temperature +<br>Population:Water +<br>Population:Temperature +<br>Water:Temperature +<br>Population:Water:Temperature | Family | 1638 |
| Population + Water + Temperature +<br>Population:Water +<br>Population:Temperature +<br>Water:Temperature | Family | 1605.2 |
| Population + Water + Temperature +<br>Population:Water +<br>Population:Temperature | Family | 1603.4 |
| Population + Water + Temperature +<br>Population:Water +<br>Water:Temperature | Family | 1577.5 |
| Population + Water + Temperature +<br>Population:Temperature +<br>Water:Temperature | Family | 1579.9 |
| Population + Water + Temperature +<br>Population:Water | Family | 1577.3 |
| Population + Water + Temperature +<br>Water:Temperature | Family | 1550.5 |
| Population + Water + Temperature | Family | 1549.3 |
| Population + Water | Family | 1541.7 |
| Population + Temperature | Family | 1546.8 |
| Water + Temperature | Family | 1551.1 |
| Population | Family | 1541.9 |
| Water | Family | 1543.9 |
| - | Family | 1541.7 |
| - | - | 1538.3 |

### Temperature and drought treatments

We imposed four treatments, a fully-factorial cross of two temperature levels and two watering levels. The temperature levels closely simulated an average growing season at the thermal extremes of the species range, which we designate as Hot and Cool treatments. Watering levels contrasted a perennial and seasonal stream, which we refer to as Well-watered and Drought treatments. A detailed description of treatments is provided in the Supplemental Materials and Methods and summarized in Fig 1. Because growth chambers cannot be subdivided, one chamber was assigned to the Hot treatment level and another to the Cool treatment level. Within each chamber, there were two Well-watered blocks and two Drought blocks. The photosynthetically active radiation in both chambers was approximately 400 µmol quanta m^−2^ s^−1^ and set to a 16:8 light:dark cycle to simulate summer growing conditions. The growth chambers did not control humidity, but because of watering and high plant transpiration rates, the relative humidity was quite high in both temperature levels (data not shown). Lower humidity would have made the drought more severe, but low soil moisture is stressful in and of itself. The total number of plants in each treatment was: *n*_cool,dry_ = 169; *n*_cool,ww_ = 174; *n*_hot,dry_ = 176; *n*_hot,ww_ = 183. Each population had 8–12 individuals per treatment level (mean = 11, median = 11).

**Figure 1:**
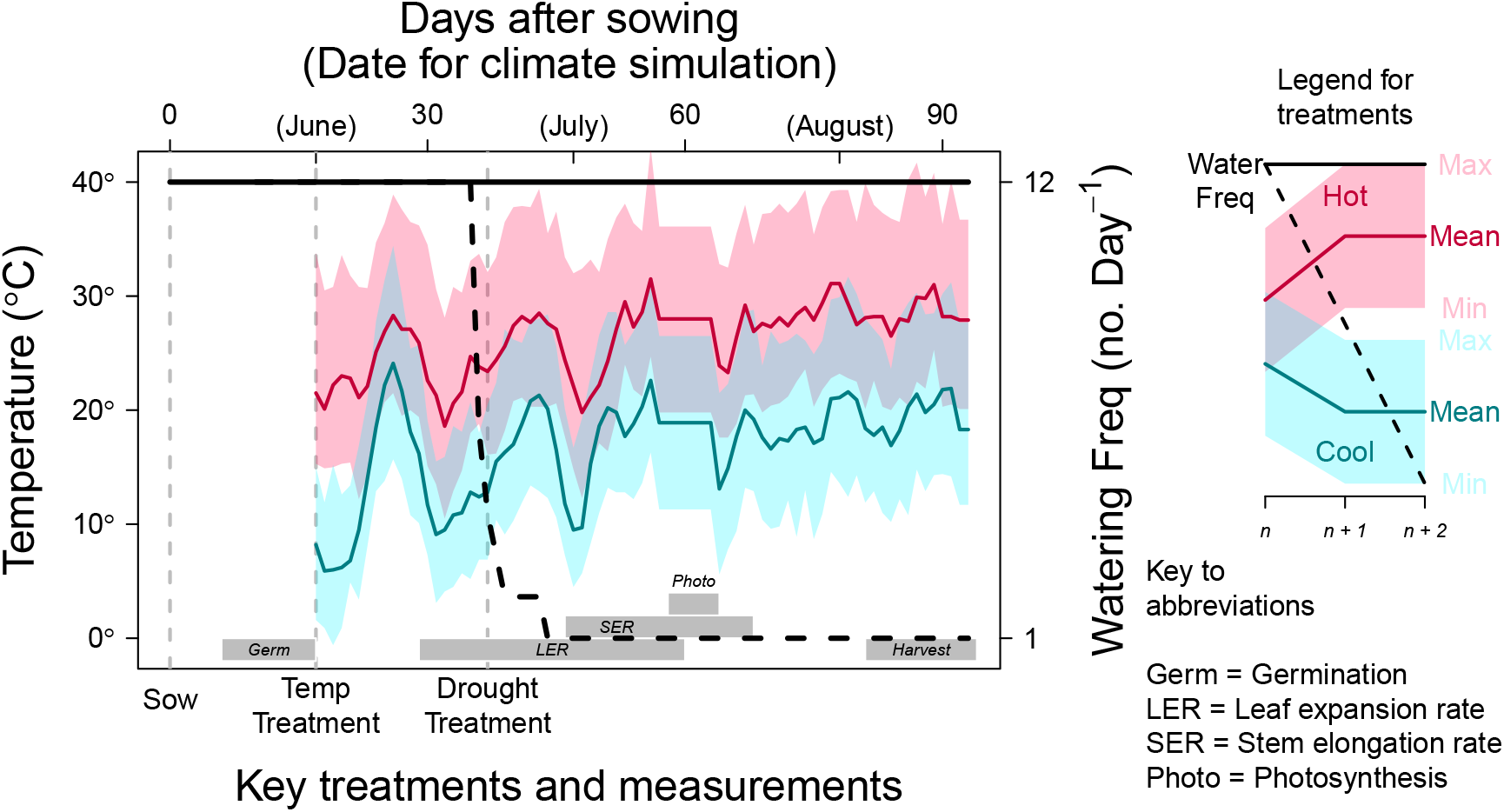
Overview of experimental treatments and timing of key trait measurements. All plants germinated within 21 days of sowing. At that time, we began temperature treatments (left axis), simulating a typical June-August weather pattern at Hot (red) and Cool (blue) sites. The bold lines track the average daily temperatures. Within each day, there was a maximum daytime temperature (top of translucent polygons) and minimum nighttime temperature (bottom of translucent polygons). The drought treatment commenced later by ramping down the frequency of bottom-watering episodes (dashed black line; right axis), while watering frequency was maintained in the control treatment (solid black line). Grey boxes on the bottom of the plot outline the period of key measurements described in the Material and Methods.

### Trait measurements

We measured five traits in response to temperature and watering treatments (Table 2).

**Table 2:** Key traits measured in this study.

| Trait | Units |
| --- | --- |
| Days to germination | day |
| Leaf expansion rate | mm day<sup>-1</sup> |
| Stem elongation rate | cm day<sup>-1</sup> |
| Photosynthetic rate | $\mu\text{mol CO}_2 \text{ m}^{-2} \text{ s}^{-1}$ |
| Mortality | probability of death |

#### Days to germination

We tested for population variation in germination rate, measured as Days to Germination, using a lognormal survival model fit using the survreg function in the R package survival version 2.38 (Therneau, 2015). We treated Population as a fixed effect and Family as random effect using a Γ frailty function. Statistical significance of the Population effect was determined using analysis of deviance. Note that, unlike other traits discussed below, we did not include Block, Treatment, or Population × Treatment interactions because during germination plants had not been placed into blocks and treatments had not yet been applied.

#### Growth rate: leaf expansion and stem elongation

We measured growth rate during two phases: leaf expansion and stem elongation. Growth measurements were taken during the early vegetative stage. We censused leaf length twice per week shortly after the emergence of true leaves from 12 May – 12 June (28–59 days after sowing), resulting in 10 measurements. We ceased measuring leaf length once it appeared to asymptote and growth shifted to stem elongation. We also censused plant height on 7 occasions (twice per week) between 29 May and 20 June (45 to 67 days after sowing) until plants began to initiate floral buds. Thus all growth measurements occured during the vegetative, prereproductive phase. Both leaf expansion and stem elongation were modelled separately as second-order polynomials. We used empirical Bayes’ estimates of growth for each individual plant from linear mixed-effects models fit with the R package **lme4** version 1.1-12 (Bates et al., 2015).

#### Photosynthesis

During the week of 10 to 16 June (57 to 63 days after sowing), we measured daytime photosynthetic rate on a subset of 329 plants evenly spread between treatments and families within populations. The youngest, fully-expanded leaf acclimated for 3 minutes to reach steady state in a 6-cm^2^ chamber of a LI-COR 6400XT Portable Photosynthesis System (LI-COR Biosciences, Lincoln, Nebraska). We made all measurements at ambient light (400 µmol m^−2^ s^−1^ of photosynthetically active radiation), atmospheric CO_2_ (400 ppm), temperature, and moderate relative humidity. All measurements were taken between 9:00 AM and 5:00 PM (3 hours after lights turned on and 5 hours before lights turned off). During this period, we suspended normal day-to-day temperature fluctuations and set daytime temperatures to the average for that period (Cool: 26.5°; Hot: 36.1°) so that all plants within a temperature level could be measured under the same conditions. We measured photosynthesis after dry down had progressed to assess differences in photosynthetic responses to drought.

#### Mortality

We assayed mortality during twice-weekly growth measurements. We analyzed the probability of surviving until the end of the experiment as a function of population, treatment, and their interactions using a Generalized Linear Mixed Model (GLMM) assuming binomially distributed errors. We included Family and Block as random effects. We assessed significance of fixed effects using Type-II Analysis of Deviance with Wald χ^2^ tests in the R package car (Fox and Weisberg, 2011).

### Genetic variation in trait means and plasticity

For all traits (Table 2) except germination (see above), we tested for Population, Treatment (Temperature, Water, and Temperature × Water), and Population × Treatment interactions (Population × Temperature, Population × Water, and Population × Temperature × Water). We interpreted significant Population effects to indicate genetic variation in trait means and Population × Treatment interactions to indicate genetic variation in plasticity. As mentioned above, we used survival and GLMM models for germination rate and mortality, respectively. For all other traits, we used mixed model ANOVAs with Family and Block included as random factors. We fit models using restricted maximum likelihood in Imer, a function in the R package **lme4** (Bates et al., 2015). We determined significant fixed effect terms using a step-wise backward elimination procedure implemented with the step function in the R package **ImerTest** version 2.0-32 (Kuznetsova et al., 2016). We used Satterthwaite’s approximation to calculate denominator degrees of freedom for *F*-tests. We also included days to germination as a covariate in growth analyses. To ensure that Population and Treatment effects were specific to a particular growth phase, we included germination day as a covariate in leaf expansion and stem elongation analyses.

Failure to detect a significant effect could be the result of Type-2 error, so we complemented step-wise ANOVA (see above) by comparing effect sizes calculated in the full model. The full model contains all main effects, two-way interactions (Population × Temperature, Population × Water), a three-way interaction (Population × Temperature × Water), and random effects. For linear mixed-effects models (leaf expansion, stem elongation, and photosynthesis) we used mean-squared error as a measure of effect size; for GLMM (mortality) we used χ^2^ as a measure of effect size. We did not include germination rate because no Population × Treatment effects were estimated. The difference in effect size of Population versus Population × Treatment is:

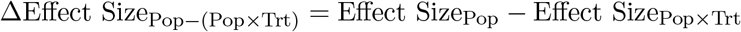

We calculated ∆Effect Size_Pop-(Pop×Trt)_ for all two-and three-way Population × Treatment interactions for each trait. To determine whether ∆Effect Size_Pop-(Pop×Trt)_ was significantly different than 0, we calculated 95% confidence intervals using 1000 parametric bootstrap samples simulated from fitted models. If the 95% confidence interval for a given ∆Effect Size_Pop-(Pop×Trt)_, was greater than zero then we concluded that the Population effect size was significantly larger than the Population × Treatment effect size, and vice versa if the confidence internval was less than zero. If the confidence spanned zero, then the effect sizes are not significantly different.

### Principal components of germination, growth, and photosynthesis

For each single-trait model above, we extracted the Population coefficient (factoring out Treatment and other effects). The multivariate distribution of these coefficients was then summarized using principal components analysis. The first principal component of these traits (TraitPC1) loaded positively with germination, growth, and photostynthetic rate, therefore we define this as a phenotypic axis delineating fast to slow growth.

### Identifying putative selective agents

Latitudinal clines are common, but it is often difficult to ascribe this variation to a particular selective agent. To reiterate, we tested three non-mutually exclusive hypotheses about how such latitudinal clines emerge: 1) one or two climatic variables explain latitudinal trait variation; 2) latitude is a proxy for multiple climatic factors that together shape trait variation; and 3) latitude integrates selection in a broader climatic neighborhood. We found that a population’s position along TraitPC1 correlated strongly with the latitude of origin (see Results) and next used Random Forest regression (Liaw and Wiener, 2002) to identify putative climatic factors underlying trait-latitude associations in *E. cardinalis*. We reasoned that if we identified a single climatic factor that explained more trait variation than latitude, then this would suggest that factor is a key selective agent underlying the latitudinal cline (Hypothesis 1). On the other hand, if multiple climatic factors together are necessary to explain trait variation, then this would suggest that many climatic factors together have imposed selection for the latitudinal cline (Hypothesis 2). We hereafter refer to factors identified in this analysis as ‘Climate-TraitPC1’ variables.

To test Hypothesis 3 about climatic neighborhoods driving selection, we directly competed local with neighborhood climate. The logic is that if the climatic analysis can identify candidate climatic factors important for local adaptation, then stronger correlations with neighbourhood climate would suggest a role for gene flow. We used the immediate collection location for local climate. For climate neighborhoods, we sampled climate at 1000 random points (at 90-m resolution) within a 62-km radius buffer around the collection and took the average. We chose this buffer radius based on population genetic structure, as inferred from ≈25,000 restriction-site associated SNPs among 49 populations from across the range (Paul et al., In review). Spatial autocorrelation in allele frequencies persists for 62 km. However radii of 10 km^2^ and 100 km^2^ resulted in similar outcomes (data not shown). Since *E. cardinalis* is found exclusively in riparian areas, we only selected points along streams using the National Hydrogeoraphy Dataset (United States Geological Survey, 2015). Climatic means and variances (see below) were weighted by their climatic suitability as determined using a multimodel ensemble average of ecological niche models (Angert et al., 2016). In addition to competing local and neighborhood climate, we compared the univariate correlation between local and neighborhood climate with TraitPC1 and Latitude using paired t-tests. We adjusted degrees of freedom to account for the fact that many climatic factors are highly correlated and not independent. Specifically, we calculated the effective number of independent climatic factors (*M*_eff_) using the formula *M*_eff_ = 1 + (M − 1)(1 − Var(λ)/*M*) (Chevrud, 2001), where *M* is the original number of climatic factors and λ are the eigenvalues of the correlation matrix of all climatic factors.

To help eliminate potentially spurious correlations between TraitPC1 and climate, we tested for overlap between climatic variables that best predict latitude of all *E. cardinalis* occurrence records (see detail below), not just the 16 focal populations. We refer to these climatic factors as ‘Climate-Latitude’ variables. The logic is that climatic factors associated with both TraitPC1 and latitude for all populations are more likely to be important selective agents than climatic factors that happen to correlate with TraitPC1 but do not covary with latitude throughout the *E. cardinalis* range. If a climatic factor is driving the latitudinal cline in TraitPC1, then we expect that climatic factor will correlate strongly with latitude of occurrence localities. Therefore, we did not consider Climate-TraitPC1 variables to be candidate selective agents unless the same or very similar variable was found in the Climate-Latitude analysis. However, we do not interpret potential selective agents identified in Climate-Latitude analyses alone, because the goal was to explain the latitudinal clines in traits, not all aspects of climate that vary with latitude.

We selected Climate-Latitude and Climate-TraitPC1 variables independently using Variable Selection Using Random Forest (VSURF) algorithm in the R package **VSURF** version 1.0.3 (Genuer et al., 2016). Random Forest regression is useful for cases like ours when the number of potential predictors is similar to or greater than the number of observations (‘high *p*, low *n*’ problem). VSURF is a multistep algorithm that progressively retains or eliminates variables based on their importance over regression trees in the forest. Variable importance is defined as the average amount a climate variable reduces mean-squared error in the predicted response (TraitPC1 or Latitude), compared to a randomly permuted dataset, across all trees in the random forest (see Genuer et al. [2015] for further detail). Hence, VSURF automatically eliminates unimportant and redundant variables based on the data without having to arbitrarily choose among colinear climate variables before the analysis. We kept only variables selected for prediction, the most stringent criterion. We visually depict how we selected climatic variables in Fig 2.

**Figure 2:**
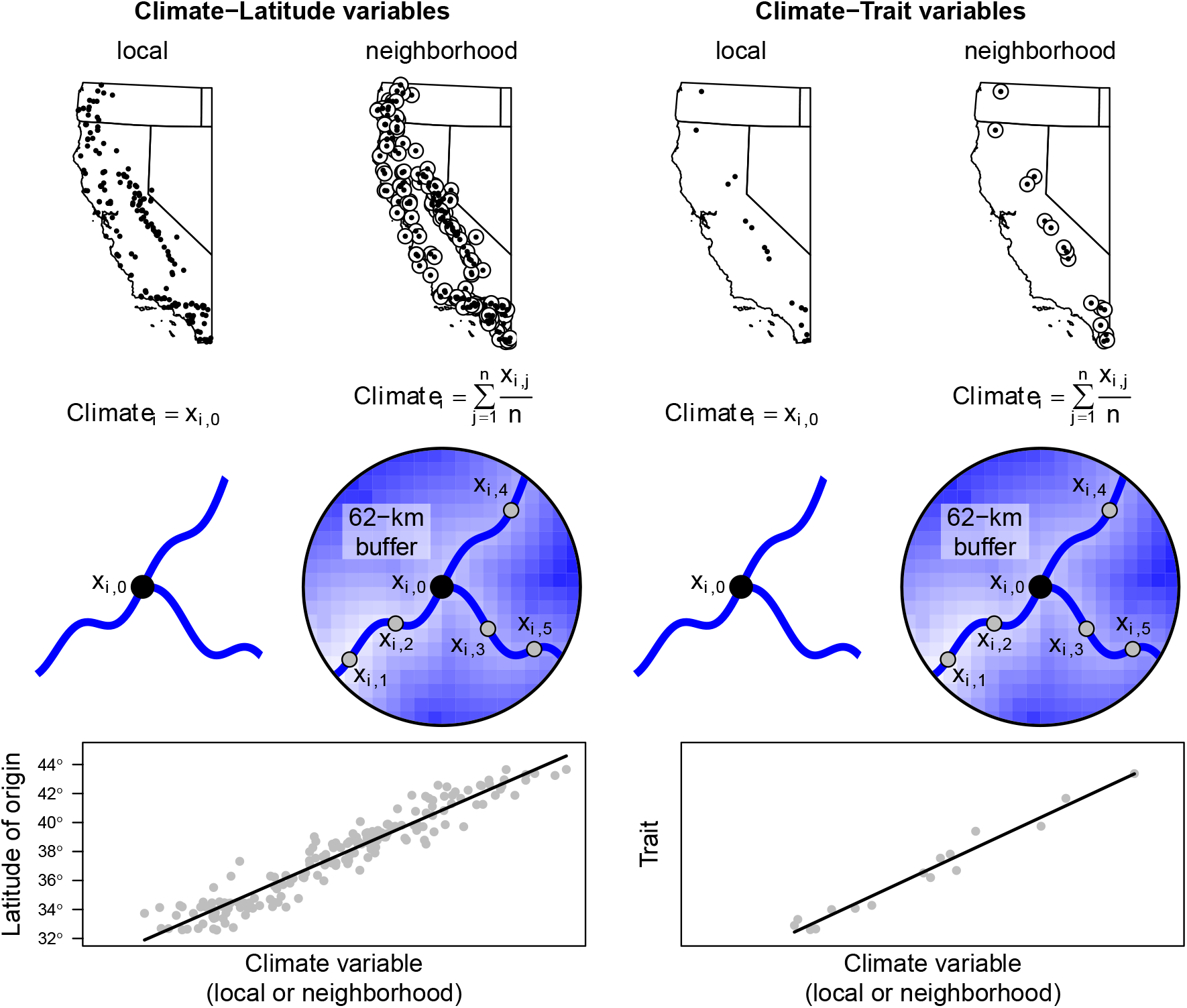
Overview of method for identifying putative climatic selective agents underlying latitudinal cline. We looked for climate variables that explained both the latitude of 356 *E. cardinalis* occurrences (‘Climate-Latitude variables’) and traits (‘Climate-Trait variables’). For Climate-Latitude variables we extracted climate data from recent occurrences located throughout California and Oregon, USA (shown in map). For Climate-Trait variables, we extracted climatic data for the 16 focal populations. For both analyses, we extracted local and neighborhood climate. Local climate refers to climate only from where a population was collected (*x_i_*,0). Neighborhood climate was calculated as the average over 1000 points in a 62-km radius climatic neighborhood (*_x_i*,1, *x_i_*,2,…), but only along stream habitats as *E. cardinalis* is riparian. We identified climatic factors that most strongly predicted latitude of occurrences (Climate-Latitude variables) and traits (Climate-Trait variables), as shown for hypothetical data in plots at the bottom of the figure.

For Climate-Latitude analyses, we compiled a representative set of 356 recent (since 2000) known *E. cardinalis* occurrences from a comprehensive set of herbarium records and an exhaustive field survey in 2010-11 (Angert et al., 2016). These occurrences were thinned by 50% to correct for uneven sampling. For both Climate-TraitPC1 analyses (16 focal populations) and Climate-Latitude (many populations), we used a 90-m digital elevation model from HydroSHEDS (Lehner et al., 2006) to extract elevation. Monthly interpolated climate layers were calculated using ClimateWNA version 5.30 (Wang et al., 2012), which accurately downscales climate data specifically for the rugged topography of western North America. For each occurence, we calculated bioclimatic variables using the biovars function in the R package **dismo** version 1.1-1 (Hijmans et al., 2016). We included 24 climatic factors, 9 from ClimateWNA and 15 bioclimatic variables (Table S2). The bioclimatic variables included all permutations of two climatic factors, temperature and precipitation, and six temporal scales (annual average, coldest quarter, warmest quarter, wettest quarter, driest quarter, or seasonality) as well as mean diurnal range, isothermality, and annual temperature range. For each variable, we calculated both a 30-year normal by averaging annual values between 1981 and 2010 and 30-year coefficient of variation, a standardized metric of interannual climatic variation. Temperatures were converted to Kelvin to be on a ratio scale appropriate for calculating the coefficient of variation (CV). In total, the VSURF algorithm selected among 96 climate variables: 24 climatic factors × 2 types (30-year average and CV) × 2 spatial scales (local and neighborhood).

**Table S2:** Climatic variables used

| Abbreviation | Climate variable |
| --- | --- |
| DD_0 | degree-days below 0°C(chilling degree-days) |
| DD5 | degree-days above 5°C(growing degree-days) |
| DD_18 | degree-days below 18°C(heating degree-days) |
| DD18 | degree-days above 18°C(cooling degree-days) |
| NFFD | number of frost-free days |
| PAS | precipitation as snow (mm) between August in previous year and July in current |
| Eref | Hargreaves reference evaporation (mm) |
| CMD | Hargreaves climatic moisture deficit (mm) |
| RH | mean annual relative humidity |
| bio1 | annual mean temperature |
| bio2 | mean diurnal range (mean of monthly (max temp - min temp)) |
| bio3 | isothermality (bio2/bio7) (* 100) |
| bio4 | temperature seasonality (standard deviation *100) |
| bio5 | max temperature of warmest month |
| bio6 | min temperature of coldest month |
| bio7 | temperature annual range (bio5-bio6) |
| bio8 | mean temperature of wettest quarter |
| bio9 | mean temperature of driest quarter |
| bio10 | mean temperature of warmest quarter |
| bio11 | mean temperature of coldest quarter |
| bio12 | annual precipitation |
| bio15 | precipitation seasonality (coefficient of variation) |
| bio16 | precipitation of wettest quarter |
| bio17 | precipitation of driest quarter |
| bio18 | precipitation of warmest quarter |
| bio19 | precipitation of coldest quarter |

## Results

### A coordinated latitudinal cline in germination, growth, and photosynthesis

There are strong genetically-based trait differences in time to germination, growth, and photosynthetic rate among populations of *E. cardinalis*, as evidenced by large and significant population effects for these traits (Table 3). A single principal component captured 71.6 % of the trait variation among populations, defining an axis of variation from fast to slow growth. A population’s position along this axis strongly covaried with its latitude of origin; southern populations grew faster than northern populations (Fig 3). There were similar latitudinal clines for individual traits underlying PC1 (Figures S1 to S4).

**Table 3:** Summary of Population, Treatment, and Population × Treatment effects. We used different statistical modeling for the diverse traits assayed – glmer: generalized linear mixed model using the R package **lme4** (Bates et al., 2015); Imer: linear mixed model using the R package **lme4** (Bates et al., 2015); survreg: survival regression using the R package survival (Therneau, 2015). Note that temperature and water treatments were imposed after germination, hence are not applicable to this trait. Complete analysis of variance/deviance tables for each trait are available in the Supporting Information. Key to statistical significance: \**P* < 0.05; ** *P* < 0.01; *** *P* < 0.001

| Trait | Germination | Leaf expansion | Stem elongation | Photosynthesis | Mortality |
| --- | --- | --- | --- | --- | --- |
| Statistical model | survreg | lmer | lmer | lmer | glmer |
| Population | *** | *** | *** | *** |  |
| Temperature | NA | *** | *** | ** | *** |
| Water | NA | * |  |  | *** |
| Pop × Temp | NA | | | * | |
| Pop × Water | NA | * | | | |
| Temp × Water | NA | | | | *** |
| Pop × Temp × Water | NA | | | | |

**Figure 3:**
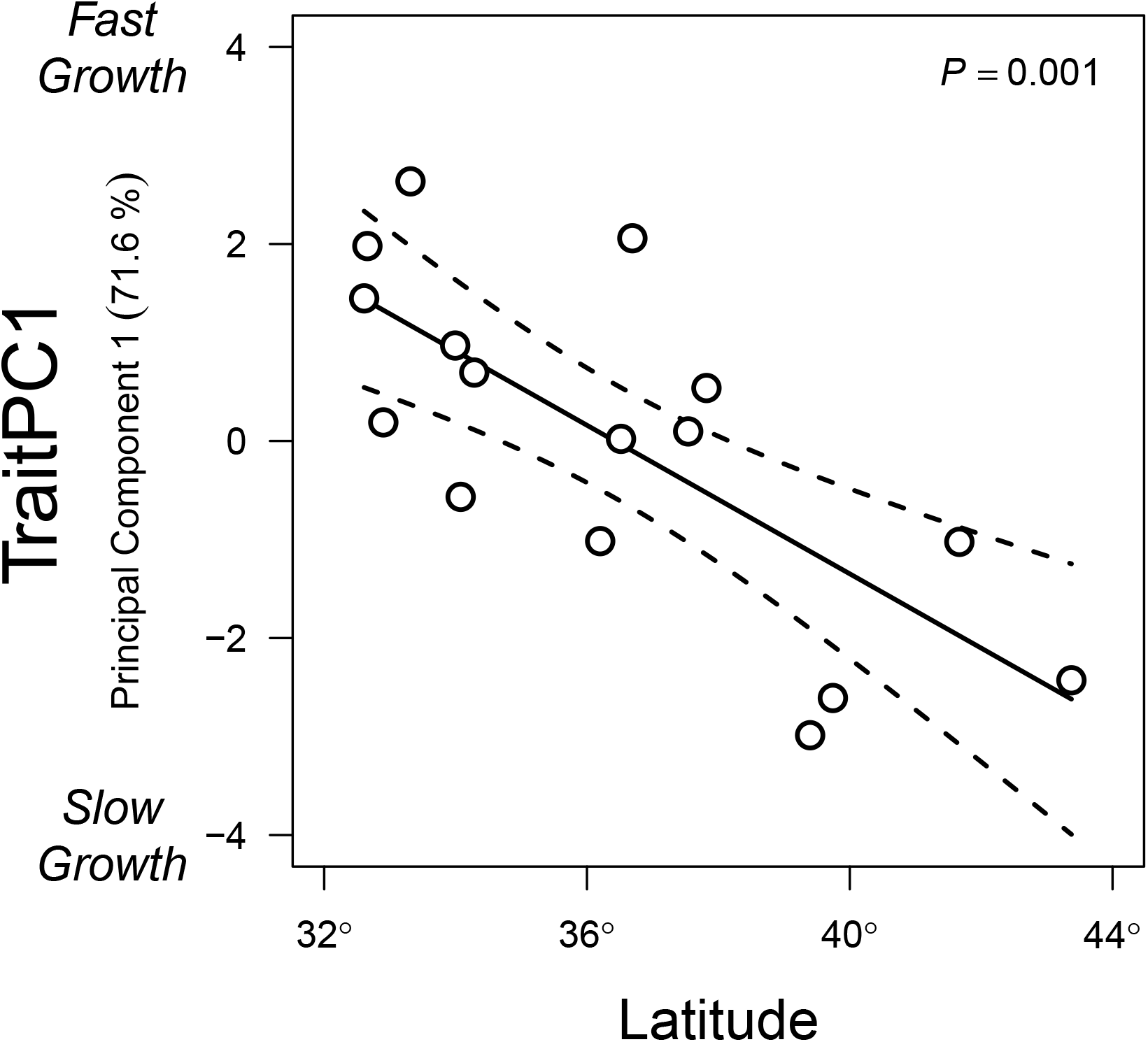
Trait variation, from fast to slow growth, is closely associated with latitude. Each point is a population’s latitude of origin (x-axis) and position along the slow to fast growth axis (y-axis), defined as Principal Component 1 of four traits (see Material and Methods). The line and 95% confidence intervals were estimated using linear regression.

### Little evidence for variation in plasticity

In contrast to the genetic variation in trait means described above, we found little evidence of G×E in *E. cardinalis*. There were only two statistically significant Population × Treatment interactions (Table 3, Fig. S5), but these were not strong compared to Population and Temperature effects. Otherwise, populations responded similarly to treatments: faster growth in the hot treatment, slower growth in the dry treatment, and high mortality in the hot, dry treatment (Table 3). Complete ANOVA tables are available in the Supporting Information (Tables S3 to S6).

The effect size of Population was significantly larger than that for Population × Treatment interactions (Fig. S6) in most cases. For leaf expansion, Population had a significantly larger effect size than Population × Treatment interactions in 2 of 3 comparisons (Fig. S6A). For stem elongation (Fig. S6B) and mortality (Fig. S6D), Population effect sizes were significantly larger than all Population × Treatment interactions. For Photosynthesis, Population and Population × Treatment effect sizes were not significantly different (Fig. S6C), presumably because we had a smaller sample size.

### Neighborhood climatic variability best explains latitudinal cline

Interannual variation in climate averaged over each populations’s climatic neighborhood correlated most strongly with trait variation and latitude of *E. cardinalis* occurrences (Fig. 4, Table S7). All 16 Climate-Latitude and 3 Climate-TraitPC1 variables were neighborhood rather than local variables (Fig. 4). In fact, neighborhood climate almost always correlated better with TraitPC1 and Latitude than local climate (Fig. 5). On average, neighborhood Climate-TraitPC1 correlation coefficients were 0.16 higher than correlations with local-scale climate variables (paired *t*-test, *t* = 7.87, d.f. = 33.6, *P* = 3.94 × 10^−9^). Likewise, neighborhood Climate-Latitude correlation coefficients were 0.13 higher than those for local-scale climate (paired *t*-test, *t* = 6.71, d.f. = 36.8, *P* = 7.22 × 10^−8^). Among Climate-Latitude and Climate-TraitPC1 variables, neighborhood climatic variability over 30 years (1981–2010) in either winter precipitation (bio16_σ_) and/or temperature (bio11_σ_) are the strongest candidates to explain the latitudinal cline in *E. cardinalis* (see Table S2 for a key to climate variable abbreviations). Note that the coefficient of variation of a climatic factor is subscripted with σ whereas the mean is subscripted with µ. More specifically, greater winter precipitation variability and lower winter temperature variability are associated with Southern latitudes and higher TraitPC1 values (Fig. 6A,B). Neighborhood interannual variation in winter precipitation (bio16_σ_) was the most important Climate-Latitude variable (Fig. 4A). However, neighborhood bio16_σ_ did not overlap with Climate-TraitPC1 variables (Fig. 4B). We nevertheless consider it a plausible candidate for two reasons. First, neighborhood bio16σ correlated strongly with TraitPC1 (Fig. 6A). Second, one of the most important Climate-TraitPC1 variables (neighborhood bio15_σ_; Fig. 6B,C) is very similar to bio16_σ_. In Mediterranean climates like California, most precipitation occurs in the wettest quarter (winter), so years with low winter precipitation also have low precipitation seasonality. Hence, highly variable year-to-year winter precipitation at lower latitude (Fig. 6D) is closely associated with large swings in precipitation seasonality (Fig. 6C).

**Figure 4:**
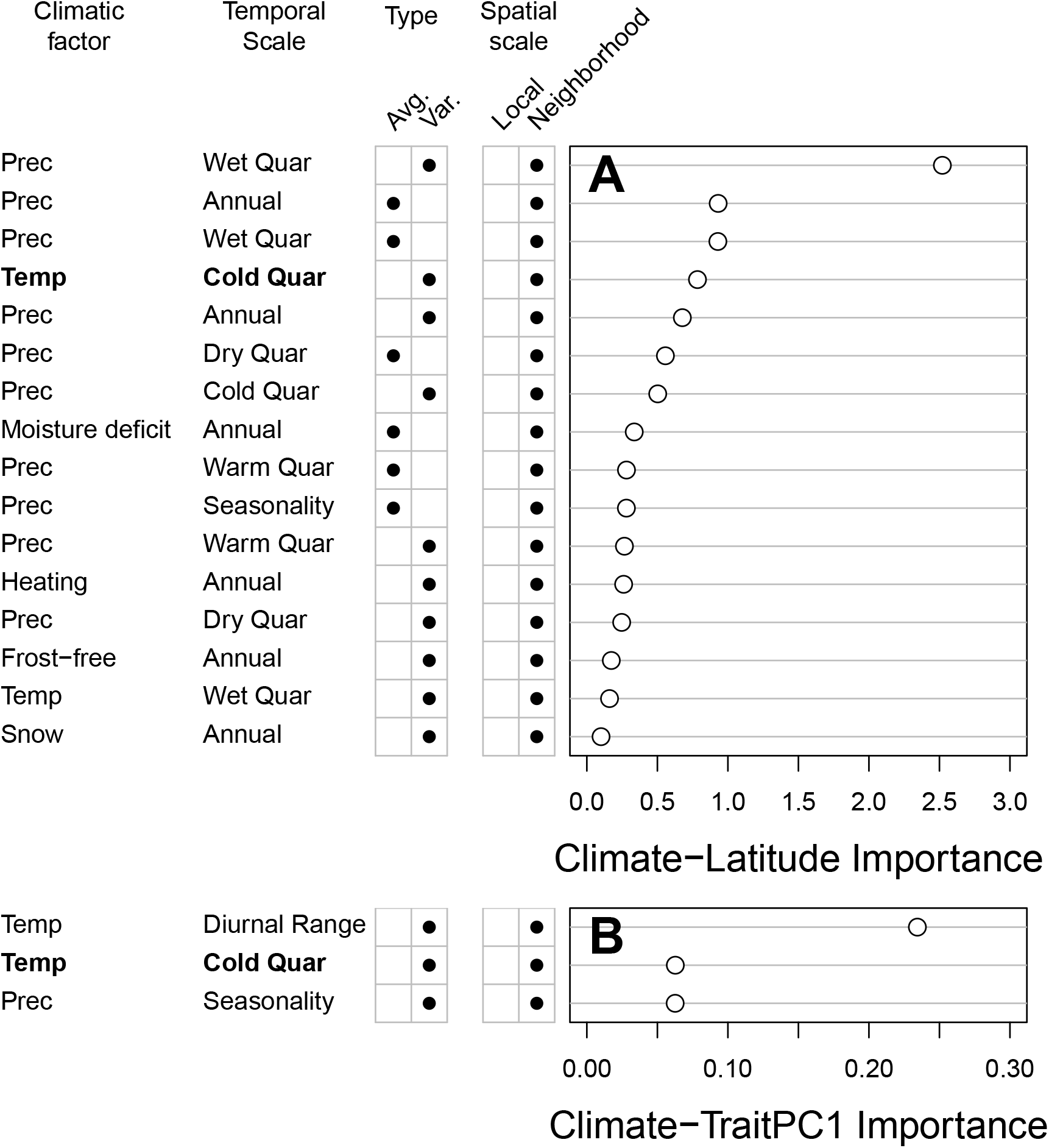
Climatic variation integrated over climatic neighborhood is closely correlated with latitude of *E. cardinalis* and trait variation. A. Using Random Forest regression, we identified 16 climatic variables significantly (high importance) associated with latitude of *E. cardinalis* occurrences. B. Only one of of the most important Climate-Latitude variables (in bold) was among the most important Climate-TraitPC1 variables. Variable importance is defined as the average amount a climate variable reduces mean-squared error in the predicted response (TraitPC1 or Latitude), compared to a randomly permuted dataset, across all trees in the random forest (see Genuer et al. [2015] for further detail). Note that the Importance values in A and B are not comparable because the dependent variables (Latitude and Trait PC1, respectively) are on different scales. Climatic variables (left of A; right of B) are defined by four qualities: Climatic factor – Temperature (Temp), Precipitation (Prec), Heating degree-days (Heating), Snow (precipitation as snow); Temporal scale – Annual, Coldest quarter (Cold Quar), Warmest Quarter (Warm Quar), Wettest quarter (Wet Quar), Driest Quarter (Dry Quar), or Seasonality; Type – 30-year average (Avg.) or coefficient of variation (Var.); Spatial scale – local or 62-km radius climatic neighborhood.

**Figure 5:**
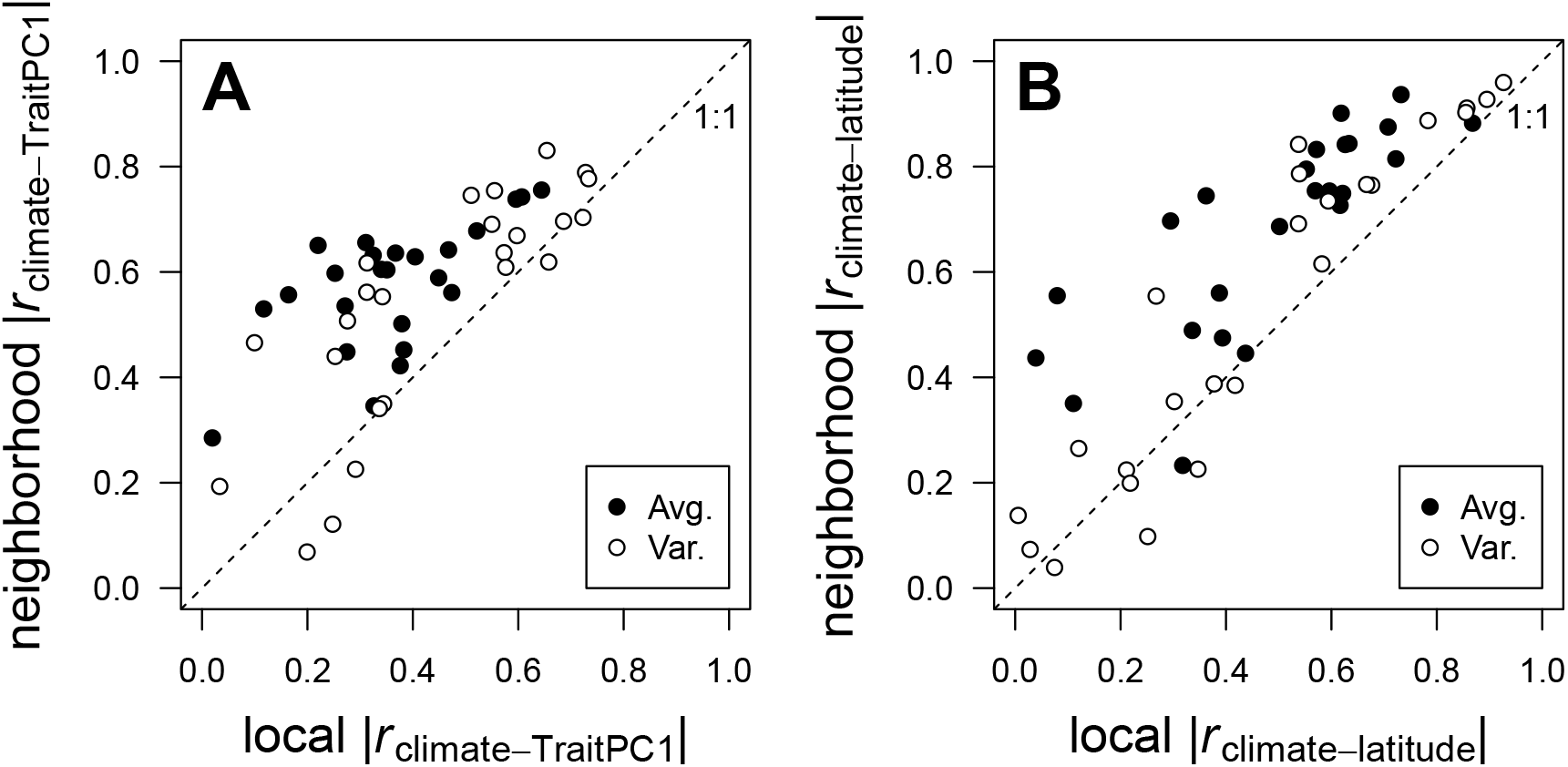
Neighborhood climate predicts TraitPC1 (‘Climate-trait’, panel A) and Latitude of occurences (‘Climate-latitude’, panel B) better than local climate. Each point is the absolute value of the Pearson correlation coefficient (|*r*|) between TraitPC1 (A) or latitude (B) for 24 climatic factors, for which we used both the 30-year mean (closed circles) and coefficient of variation (open circles). Most points lie above the 1:1 line, indicating stronger correlations with neighborhood compared to local climate. Neighborhood climate was integrated over a 62-km radius around focal populations (see text for further detail).

**Figure 6:**
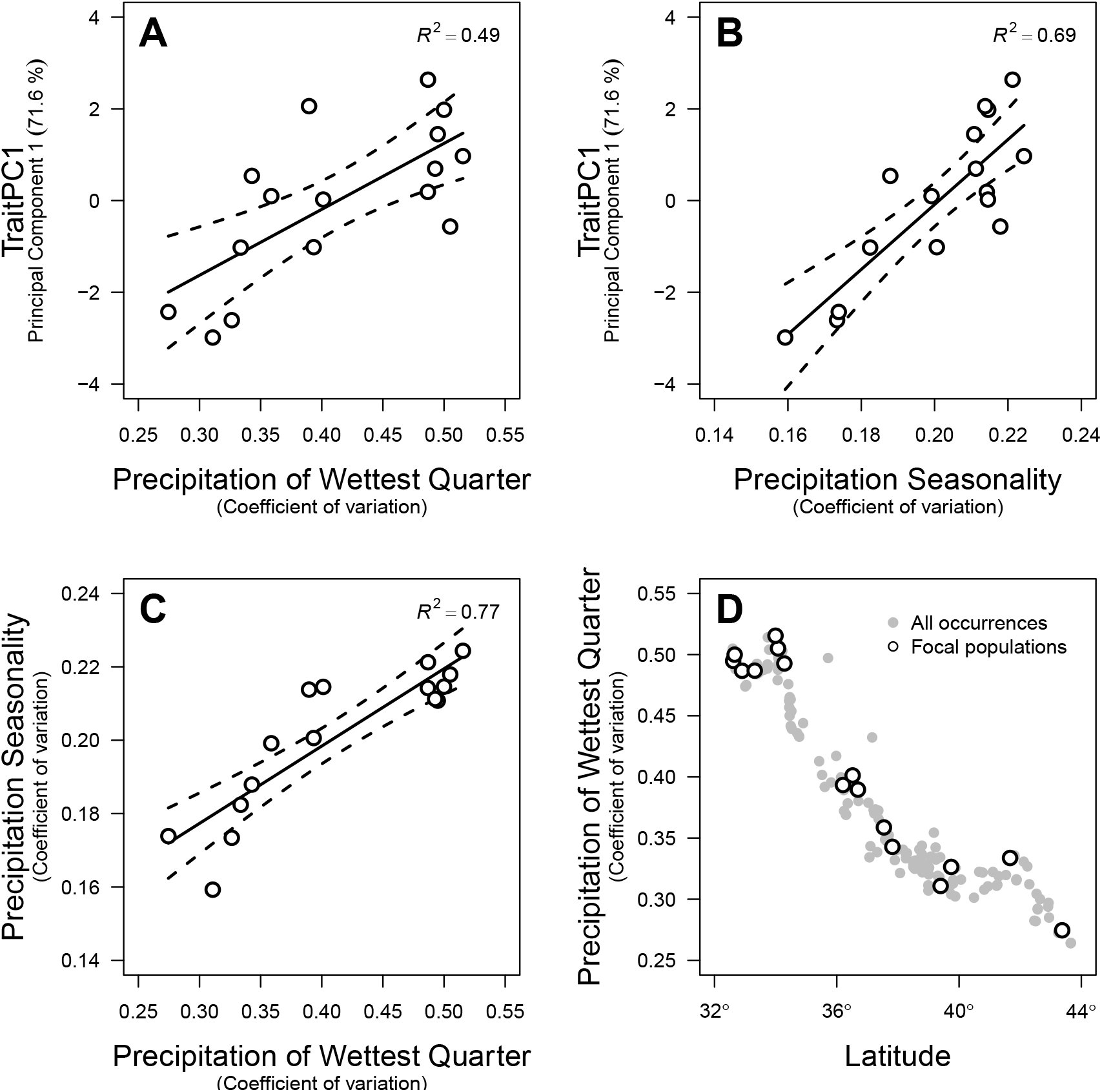
Variation in precipitation is correlated with TraitPC1 and latitude. A. Greater values of TraitPC1 are associated with greater interannual variation in precipitation of the wettest quarter. This was the most important Climate-Latitude variable, but not among the most important Climate-TraitPC1 variables. B. However, a closely related parameter, interannaul variation in precipitation seasonality, was among the most important Climate-TraitPC1 variables. C. Across focal populations, variation in precipitation of the wettest quarter and seasonality are closely correlated. D. Southern populations of *E. cardinalis* experience much greater interannual vari-ationi in precipitation. In all panels, we report climatic neighborhood values (see Material and Methods). Regression lines, 95% confidence intervals, and coefficients of determination (*R*^2^) were calculated using linear regression.

Interannual variation in temperature of the coldest quarter (neighborhood bio11_σ_) is another plausible candidate because it was the only variable in both Climate-Latitude and Climate-TraitPC1 analyses (Fig. 4). Neighborhood bio11_σ_ explained more variation in TraitPC1 than latitude (latitude r^2^ = 0.55 vs. bio11_σ_ r^2^ = 0.6; Fig. S7), whereas neighborhood bio16_σ_ did slightly worse (bio16_σ_ r^2^ = 0.49). Models using bio15_σ_ or bio11_σ_ to predict TraitPC1 also had significantly lower Akaike Information Criteria (AIC) than the latitude model (AIC of different models – bio15_σ_: 48.5; bio11_σ_: 52.4; latitude: 54.5). The best two-factor model including both neighborhood bio15_σ_ and bio11_σ_ did not significantly improve explanatory power (r^2^ = 0.71, AIC= 49.2). In summary, either variation in precipitation or temperature seasonality may be important selective agents, but there is no strong evidence that they are both important. The most important Climate-TraitPC1 variable, neighborhood variation in mean diurnal range (bio2_σ_; Fig. 4B) did not have any obvious similarity to Climate-Latitude variables. Given the large number of potential associations, we therefore think this may be a spuriously strong relationship.

## Discussion

We found evidence for one of two common signatures of local adaptation in the perennial herb *Erythranthe cardinalis*. Latitudinal clines in germination rate, photosynthesis, and growth suggest adaptive differentiation in important physiological traits of the species. However, we caution that these are candidate adaptive traits and that we cannot yet rule out nonadaptive demographic processes such as a recent range expansion toward higher latitude (Paul et al., In review; Sheth and Angert, 2017). In contrast, we found little evidence for variation in plasticity to temperature or drought. Due to low replication within families, we did not have power to assess within-population genotype-by-environment interactions, which may be present. As we discuss below, low variation in plasticity among populations may indicate that some dimensions of the fundamental abiotic niche are relatively conserved. Note that statistical power to detect significant plasticity is lower than that for differences in trait means, but the effect size of variation in plasticity was significantly less than that for trait means in most cases (Fig. S6). Finally, our results suggest that neighborhood-scale climate and interannual variation are more important selective agents than local averages. In the paragraphs that follow, we tie these results into the broader threads of evolutionary theory that might help explain why variation in physiological trait means changes clinally, whereas plastic responses to temperature and drought are relatively static. One caveat to bear in mind is that we are limited by the size of the climate grid (≈ 90 m^2^) and therefore unable to detect very fine-scale local adaptation.

Evolutionary theory indicates that the shape of fitness tradeoffs, demography, and gene flow can constrain adaptation (Levins, 1968; Ronce and Kirkpatrick, 2001; Lenormand, 2002) and hence the type of variation maintained within species. Specifically, adaptive variation can be maintained by spatially varying selection if tradeoffs are not too strong, demography is symmetric, and/or maladaptive gene flow is low. Strong tradeoffs can prevent local adaptation in spatially variable environments because selection favors habitat specialists that track a specific habitat regardless of its frequency in the environment (Levins, 1968). For example, a riparian specialist may experience similar selection in rivers of high rainfall regions and deserts, even though the habitat is much rarer in the latter. In *E. cardinalis* we found substantial genetically based variation among populations along a phenotypic axis from fast to slow growth that varied over a large spatial scale (Fig. 3). If this variation is adaptive, it suggests one of several possibilities to investigate in the future: the fitness tradeoff between low versus high latitude environments is not too strong nor swamped by demographic asymmetry or maladaptive gene flow. That is, alleles favoured at one latitude are not strongly selected against when they flow to another population, allowing locally adaptive genetic variation to be maintained by spatially heterogeneous selection. We also know from previous work that population size does not vary strongly with latitude (Angert, unpub. data). Gene flow appears to be high, but attenuates at broad spatial scales, especially between southern (< 35°N) and northern portions of the range (Paul et al., In review).

Nevertheless, local gene flow from similar environments may shape how selection varies with latitude. Theory predicts that populations will not be perfectly adapted to their immediate habitat when there is gene flow from surrounding populations with different optima (Lenormand, 2002). With spatial heterogeneity and gene flow, traits will not covary perfectly with the local optimum (Slatkin, 1978; Paul et al., 2011; Hadfield, 2016), but should instead better match the average environment experienced by nearby populations connected through gene flow, which we refer to as the climatic neighborhood. Gene flow and spatial heterogeneity may therefore be important in maintaining genetic variation (Yeaman and Jarvis, 2006). As this hypothesis predicts, climatic neighborhoods (62-km buffer around populations) correlated with traits and latitude of occurrences better than local climate (Fig. 4). We interpret this as suggestive evidence that gene flow between neighboring *E. cardinalis* populations shapes selection – populations are locally adapted to prevailing climate in their neighborhood, but perhaps not perfectly adapted to their local climate. This may not greatly constrain local adaptation because local and neighborhood climate values were generally similar in *E. cardinalis* populations (Fig. 5), at least at the resolution of ClimateWNA (90 m^2^). Therefore, we would predict in reciprocal transplants that populations whose local climate is farther from their neighborhood average would be less well adapted than those close to their neighborhood average.

It is reasonable to predict that southern populations, which appear to experience more frequent drought years (see below), might have physiological adaptations to respond to drought stress to survive and grow in drier soil. We found little evidence for this type of drought tolerance; all populations responded to drought and temperature similarly (Table 3). Plants grew faster in the Hot treatment, but there was little effect of drought on growth. Rather, the effects of drought took longer to materialize but resulted in high mortality, especially in the Hot treatment. However, there was no differential mortality among populations in this treatment. Although our results indicate that this axis of the species niche may be constrained, plants have multiple ways to resist drought through both tolerance and escape (Ludlow, 1989; Kooyers, 2015). Next, we consider why drought tolerance may less important in local adaptation than a form of escape for this species.

We hypothesize that tolerance to dry soil may be constrained by a combination of strong fitness tradeoffs, demographic asymmetry, and gene flow. Soil moisture in riparian habitats where *E. cardinalis* lives is highly heterogeneous at very small spatial scales (several meters). Plants in the stream never have to tolerate drought whereas plants only a few meters away may experience extreme drought since there is little direct precipitation during the growing season in Mediterranean climates of western North America. We hypothesize that alleles confering greater drought tolerance may be quite costly in well-watered soils, and vice versa, leading to strong fitness tradeoffs. Such tradeoffs would promote specialization to one soil moisture or another, thereby inhibiting the evolution of broad environmental tolerance within a population. Demography and gene flow may reinforce niche conservatism. A new mutant with increased drought tolerance that could survive at the resource-poor margin of a population would likely be demographically overwhelmed by the larger census populations that can be maintained in higher-resource environments. Infrequent wet years may also produce most seeds, so selection is weighted towards alleles that have high fitness in the wet environment, even if dry years are more frequent (Templeton and Levin, 1979; Brown and Venable, 1986). However, demographic asymmetry should equally hinder the evolution of both drought tolerance and escape, so it should not explain why one mechanism evolves but not the other. Finally, gene flow, which is generally high among *E. cardinalis* populations within the same ecoregion (Paul et al., In review), will thwart local adaptation and reinforce specialization. Thus, the spatial grain of the environment, demographic asymmetry, and gene flow may conspire to constrain local adaptation along this environmental axis. Consistent with this hypothesis, recent record-setting droughts have caused the decline or even local extinction of some natural populations of *E. cardinalis* (Sheth and Angert, 2017).

In sum, these results indicate that genetic differences in physiology and growth are better candidates than plastic responses to temperature and drought as mediators of local adaptation to climate in *E. cardinalis*. Next, we would like to understand why variation in these particular traits may be adaptive. We argue that temporally more variable environments, as experienced by southern populations, select for a more ‘annualized’ life-history strategy, a form of drought escape. Demographic observations in natural populations of *E. cardinalis* reveal that southern populations tend to flower earlier at a smaller size, while northern populations invest more in vegetative growth (Sheth and Angert, 2017). In this experiment, the fastest growing plants began producing flowers in ~ 60 days (data not shown), suggesting that rapid vegetative develop may likewise affect flowering time. The association between position along the ‘fast-slow’ continuum and associated traits in *E. cardinalis* is similar to interspecific relationships between growth, functional traits, and life history (Adler et al., 2014; Salguero-Gómez et al., 2016). However, we cannot exclude unexplored factors (e.g. edaphic conditions, competitors, pollinators, etc.) which may also contribute to the latitudinal cline.

Greater investment in aboveground growth, as opposed to belowground storage for future seasons, may be favoured in climates with more frequent drought years, but maladaptive in climates with more consistent precipitation. In a stable environment where winter survivorship is assured in most years, failure to store resources may reduce lifetime fitness. But for perennial herbs in Mediterranean climates, a dry winter (rainy season) can kill the rhizomes (underground stems that store nutrients for future growth) before emergence or aboveground stems before flowering. If drought years occur frequently enough, selection may favour the fast-growing strategy because there is no advantage to storage if drought kills plants before flowering. Considering life-history strategy as a continuum from no storage (annual) to lots of storage (perennial), we hypothesize that the optimal allocation to aboveground growth is more ‘annualized’ in southern climates that have greater interannual variation in precipitation.This is a form of drought escape in that plants are investing more reproduction in the present to avoid possible drought in subsequent years, but is distinct from classic drought escape syndromes in which plants speed up development early in the season before the onset of drought.

The hypothesis that greater precipitation variability selects for an annualized life history is tentative, but consistent with theory and data from other species. Life history theory shows that less variable environments are one factor that favours the evolution of perenniality (Stearns, 1976; Iwasa and Cohen, 1989; Friedman and Rubin, 2015). Populations of the perennial *Plantago asiatica* show a similar latitudinal cline in growth and allocation to storage (Sawada et al., 1994) but attribute the cline to variation in growing season length. There are also life history clines in the closely related species *E. guttata*, but the underlying traits and climatic drivers are quite different. Annual *E. guttata* flower sooner and produce fewer stolons in response to climates with shorter seasons and more intense summer drought (Lowry and Willis, 2010; Friedman et al., 2015; Kooyers et al., 2015). In contrast, there are no truly annual (monocarpic and semelparous) populations of *E. cardinalis*. Rather, our hypothesis states that climatic variability selects on quantitative variation in allocation to growth versus storage.

In summary, we found evidence for a coordinated latitudinal cline in germination rate, photosynthesis, and growth, suggesting local adaptation. We therefore predict to find different optima for these traits in different climates. We did not find evidence that the relative performance of populations shifts with temperature or watering regime, suggesting relatively little variation in plasticity. Exploratory analysis implicate that more variable precipitation regimes at lower latitude could drive much of the latitudinal cline, though other climatic factors could also contribute. Interestingly, the climatic neighborhood may shape selective pressures more than local climate. In the future, we will use field experiments to test whether greater variation in precipitation selects for faster growth and if selection on temperature/drought responses does not vary among populations. By doing so, we aim to understand why certain physiological and developmental mechanisms, but not others, contribute to local adaptation.

## Acknowledgements

Erin Warkman and Lisa Lin helped collect data. CDM was supported by a Biodiversity Postdoctoral Fellowship funded by the NSERC CREATE program. ALA was supported by an NSERC Discovery Grant and a grant from the National Science Foundation (DEB 0950171). Four anonymous referees provided constructive comments on earlier versions of this manuscript.

## Supporting Information

### Supporting Tables

**Table S3:** Analysis of varianace (ANOVA) table on leaf expansion rate (LER) using ImerTest (Kuznetsova et al., 2016). Family and Block were included as random effects. Abbreviations: SS = sum of squares; MS = mean sum of squares (SS / df1); df1 = numerator degrees of freedom; df2 = denominator degrees of freedom.

|  | SS | MS | df1 | df2 | *F-value* | *P-value* |
| --- | --- | --- | --- | --- | --- | --- |
| Day to Germination | 12.12 | 12.12 | 1 | 637 | 35.21 | $4.9 \times 10^{-9}$ |
| Population | 22.22 | 1.48 | 15 | 118 | 4.3 | $2.5 \times 10^{-6}$ |
| Temperature | 80.42 | 80.42 | 1 | 5 | 233.61 | $2.6 \times 10^{-5}$ |
| Water | 4.1 | 4.1 | 1 | 5 | 11.92 | 0.019 |
| Temperature × Water | 0.03 | 0.03 | 1 | 4 | 0.07 | 0.801 |
| Population × Temperature | 2.76 | 0.18 | 15 | 547 | 0.53 | 0.925 |
| Population × Water | 9.66 | 0.64 | 15 | 562 | 1.87 | 0.024 |
| Population × Temperature × Water | 4.11 | 0.27 | 15 | 530 | 0.78 | 0.700 |

**Table S4:** Analysis of varianace (ANOVA) table on stem elongation rate (SER) using ImerTest (Kuznetsova et al., 2016). Family and Block were included as random effects. Abbreviations: SS = sum of squares; MS = mean sum of squares (SS / df1); df1 = numerator degrees of freedom; df2 = denominator degrees of freedom.

|  | SS | MS | df1 | df2 | *F-value* | *P-value* |
| --- | --- | --- | --- | --- | --- | --- |
| Day to Germination | 3.6 | 3.6 | 1 | 662 | 21.1 | $5.1 \times 10^{-6}$ |
| Population | 12 | 0.8 | 15 | 113 | 4.7 | $5.8 \times 10^{-7}$ |
| Temperature | 12.4 | 12.4 | 1 | 6 | 72.8 | $1.5 \times 10^{-4}$ |
| Water | 0.6 | 0.6 | 1 | 5 | 3.7 | 0.113 |
| Temperature × Water | 0.9 | 0.9 | 1 | 4 | 5.2 | 0.093 |
| Population × Temperature | 3.6 | 0.2 | 15 | 549 | 1.4 | 0.126 |
| Population × Water | 2.8 | 0.2 | 15 | 536 | 1.1 | 0.330 |
| Population × Temperature × Water | 1.5 | 0.1 | 15 | 518 | 0.6 | 0.874 |

**Table S5:** Analysis of varianace (ANOVA) table on photosynthetic rate using **ImerTest** (Kuznetsova et al., 2016). Family and Block were included as random effects. Abbreviations: SS = sum of squares; MS = mean sum of squares (SS / df1); df1 = numerator degrees of freedom; df2 = denominator degrees of freedom.

|  | SS | MS | df1 | df2 | *F-value* | *P-value* |
| --- | --- | --- | --- | --- | --- | --- |
| Population | 347.7 | 23.2 | 15 | 78 | 3.02 | $7.5 \times 10^{-4}$ |
| Temperature | 134.1 | 134.1 | 1 | 6 | 17.46 | $6.4 \times 10^{-3}$ |
| Water | 51 | 51 | 1 | 4 | 6.64 | 0.066 |
| Temperature × Water | 0.7 | 0.7 | 1 | 3 | 0.09 | 0.781 |
| Population × Temperature | 218.6 | 14.6 | 15 | 263 | 1.9 | 0.024 |
| Population × Water | 87.7 | 5.8 | 15 | 233 | 0.76 | 0.724 |
| Population × Temperature × Water | 91.4 | 6.1 | 15 | 208 | 0.79 | 0.686 |

**Table S6:** Analysis of deviance table on the probability of mortality by the end of the experiment using Type-II Wald χ^2^ tests in the R package car (Fox and Weisberg, 2011). Family and Block were included as random effects. Abbreviations: df = degrees of freedom

| | $\chi^2$ | df | *P-value* |
| --- | --- | --- | --- |
| Population | 32 | 31 | 0.419 |
| Temperature | 31.8 | 6 | $1.8 \times 10^{-5}$ |
| Water | 69.2 | 12 | $4.6 \times 10^{-10}$ |
| Temperature × Water | 20.7 | 1 | $5.3 \times 10^{-6}$ |
| Population × Temperature | 5.6 | 15 | 0.985 |
| Population × Water | 8.6 | 15 | 0.897 |
| Population × Temperature × Water | 0.2 | 15 | 1.000 |

**Table S7:** Important climatic variables predicting latitude of *E. cardinalis* populations (‘Climate-Latitude’) and the first principal component of traits measured in a common garden (‘Climate-TraitPC1’). Local climatic variables were measured from the exact location of collection; neighborhood climatic variables were averaged from a 62-km neighborhood around population (see Material and Methods). Importance and significance were determined using the variable selection using random forests (VSURF) algorithm (see Material and Methods). Climatic variables are described in Table S2. *µ* signifies the mean of the climate variables from 1981–2010; *σ* indicates coeffiecient of variation among years.

| Climate-Latitude variables | Climate-TraitPC1 variables |
| --- | --- |
| Precipitation of wettest quarter ($\sigma$, neighborhood) | Mean diurnal range ($\sigma$, neighborhood) |
| Annual precipitation ($\mu$, neighborhood) | Mean temperature of coldest quarter ($\sigma$, neighborhood) |
| Precipitation of wettest quarter ($\mu$, neighborhood) | Precipitation seasonality ($\sigma$, neighborhood) |
| Mean temperature of coldest quarter ($\sigma$, neighborhood) | |
| Annual precipitation ($\sigma$, neighborhood) | |
| Precipitation of driest quarter ($\mu$, neighborhood) | |
| Precipitation of coldest quarter ($\sigma$, neighborhood) | |
| Hargreaves climatic moisture deficit ($\mu$, neighborhood) | |
| Precipitation of warmest quarter ($\mu$, neighborhood) | |
| Precipitation seasonality ($\mu$, neighborhood) | |
| Precipitation of warmest quarter ($\sigma$, neighborhood) | |
| Heating degree-days ($\sigma$, neighborhood) | |
| Precipitation of driest quarter ($\sigma$, neighborhood) | |
| Number of frost-free days ($\sigma$, neighborhood) | |
| Mean temperature of wettest quarter ($\sigma$, neighborhood) | |
| Precipitation as snow ($\sigma$, neighborhood) | |

**Figure S1:**
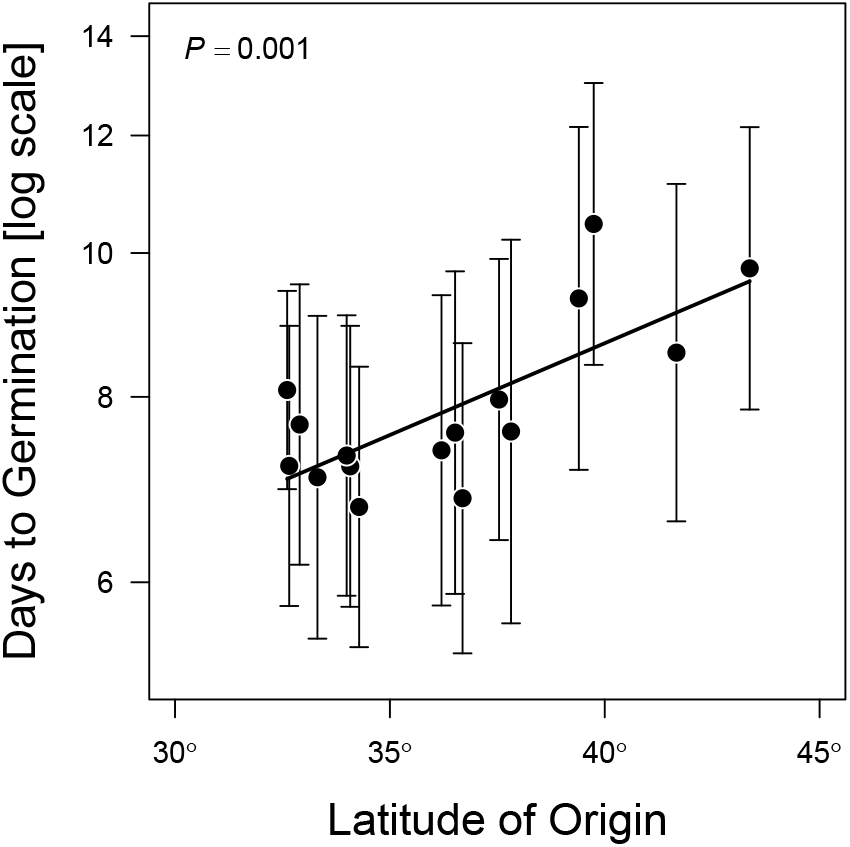
Southern populations germinate faster. Each point is a population of *E. cardinalis* showing its latitude of origin (x-axis) and model-predicted days to germination in days under growth chamber conditions (see Material and Methods). Bars around each point are 95% confidence intervals. Predicted time to germination and confidence intervals are based on survival regression (see Materials and Materials). The line is the linear regression of log(model-predicated days to germination) ~ latitude. The *P*-value of the regression is given in the upper left corner.

**Figure S2:**
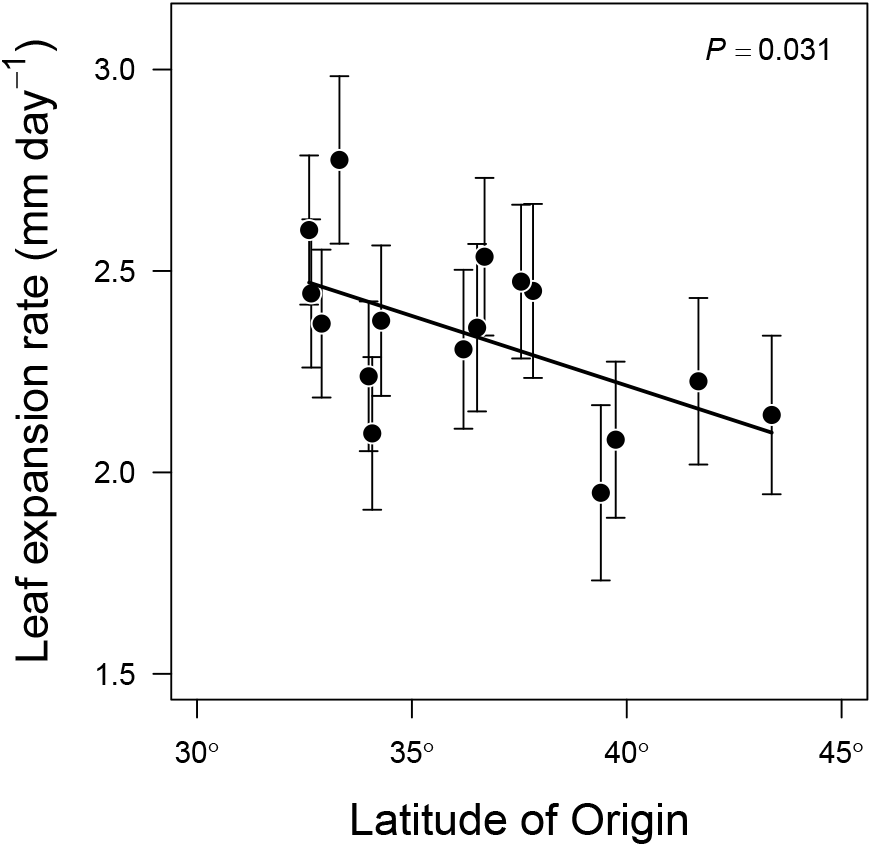
Southern populations grow faster. Each point is a population of *E. cardinalis* showing its latitude of origin (x-axis) and model-predicted leaf expansion rate during the rosette phase. Bars around each point are 95% confidence intervals. Predicted leaf expansion rate based least-square mean estimates and confidence intervals were calculated from linear mixed-effects models (see Materials and Materials). The line is the linear regression of model-predicated leaf expansion rate ~ latitude. The *P*-value of the regression is given in the upper right corner.

**Figure S3:**
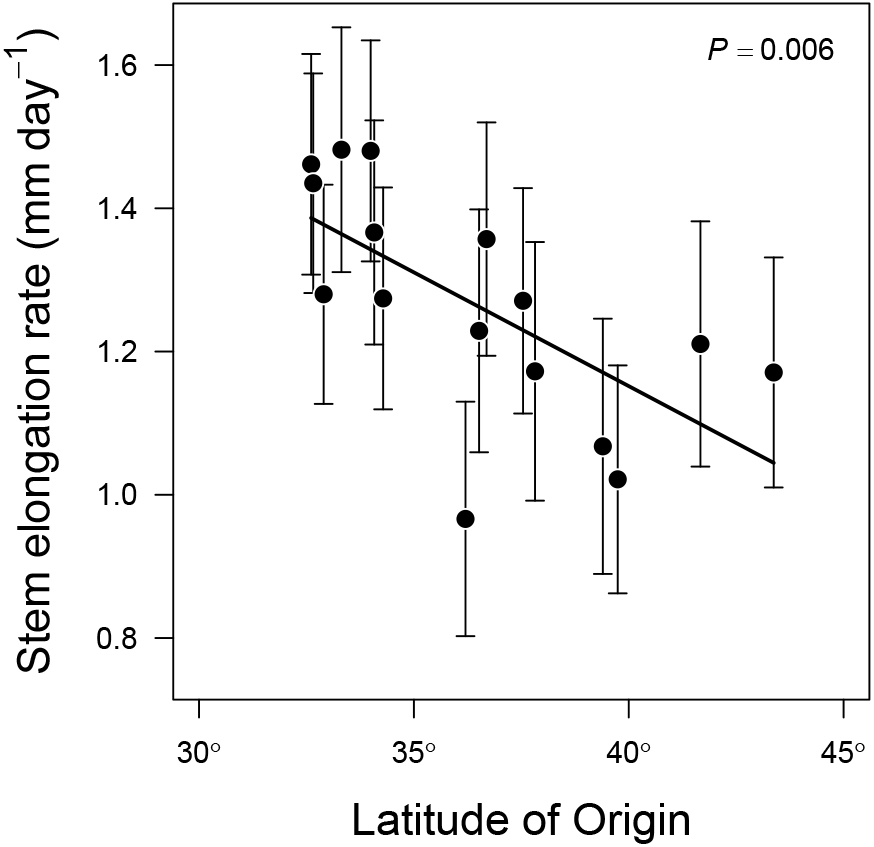
Southern populations grow faster. Each point is a population of *E. cardinalis* showing its latitude of origin (x-axis) and model-predicted stem elongation rate. Bars around each point are 95% confidence intervals. Predicted stem elongation rate based least-square mean estimates and confidence intervals were calculated from linear mixed-effects models (see Materials and Materials). The line is the linear regression of model-predicated stem elongation rate ~ latitude. The *P*-value of the regression is given in the upper right corner.

**Figure S4:**
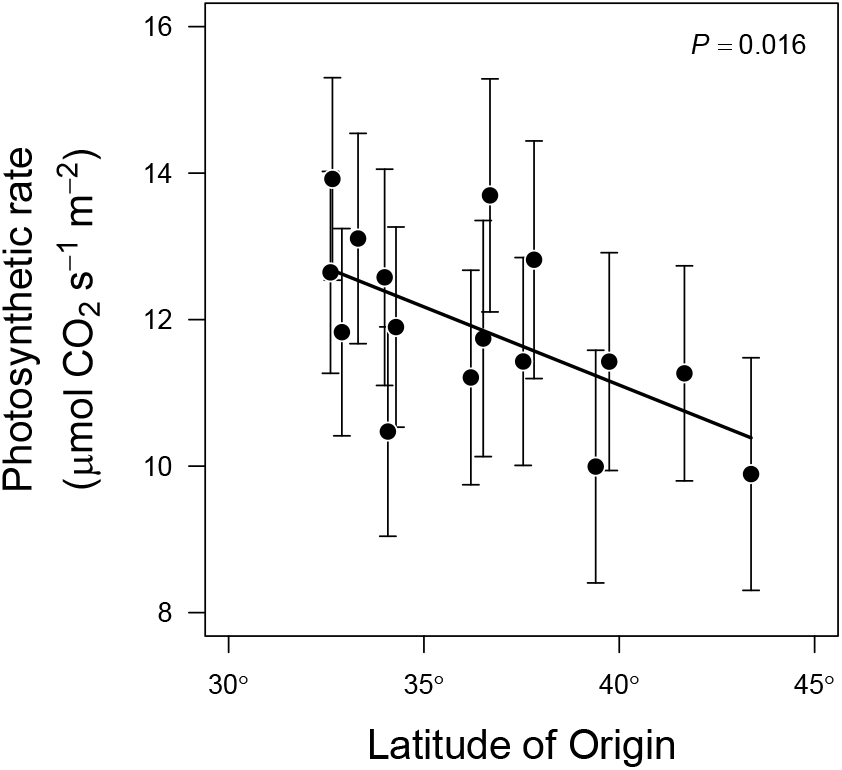
Southern populations photosynthesize faster. Each point is a population of *E. cardinalis* showing its latitude of origin (x-axis) and model-predicted instantaneous photosynthetic rate. Bars around each point are 95% confidence intervals. Predicted photosynthetic rates based least-square mean estimates and confidence intervals were calculated from linear mixed-effects models (see Materials and Materials). The line is the linear regression of model-predicated photosynthetic rate ~ latitude. The *P*-value of the regression is given in the upper right corner.

**Figure S5:**
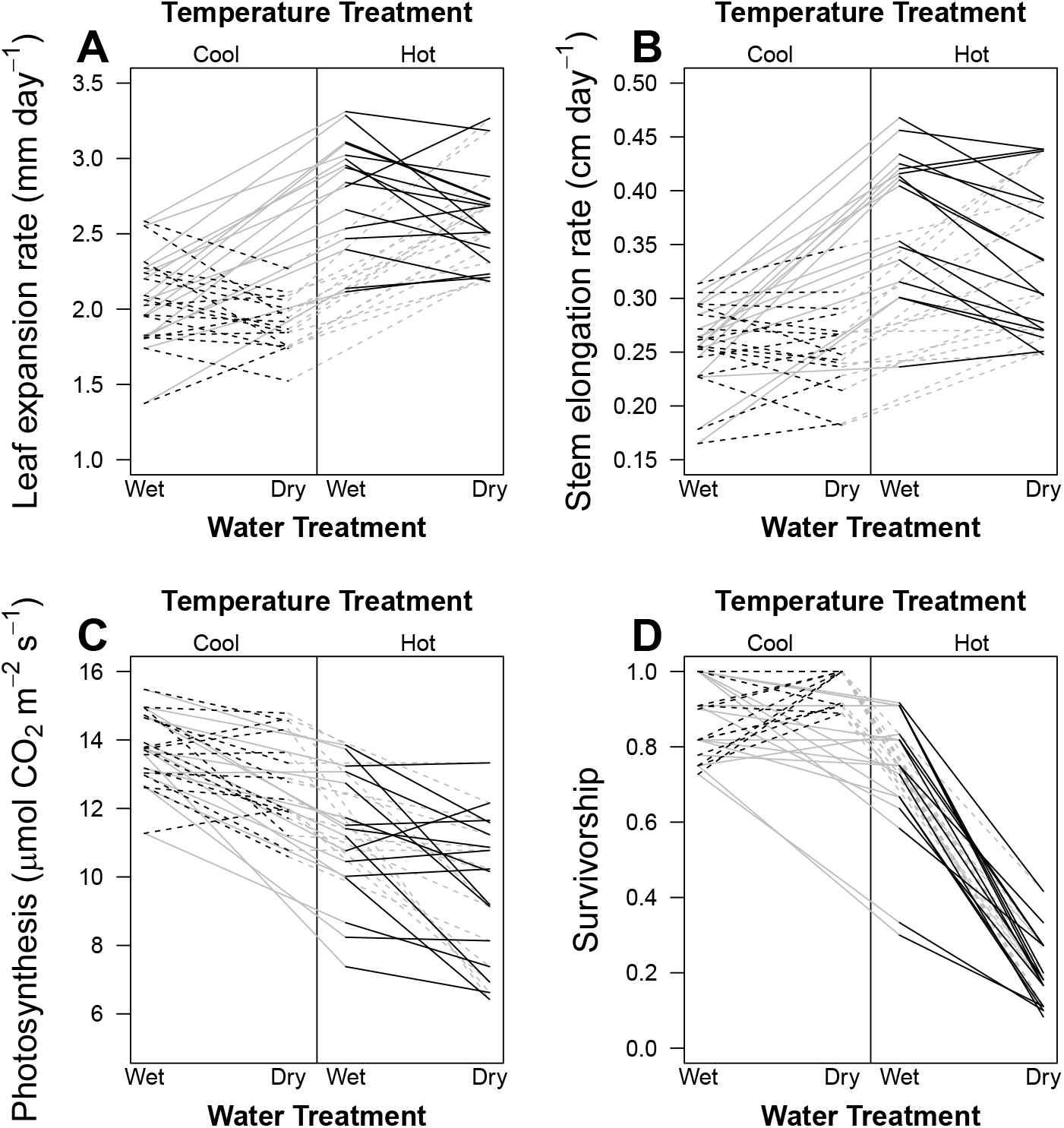
Reaction norms signify little Population × Treatment interactions. For all panels, black lines represent population-level reaction norms from Wet to Dry in the Cool temperature treatment (dashed black lines) and Hot temperature treatment (solid black lines); gray lines represent reaction norms from Cool to Hot in the Wet treatment (solid gray lines) and Dry treatment (dashed gray lines). The responses shown are (A) leaf expansion rate, (B) stem elongation rate, (C) photosynthesis, and (D) survivorship (= 1 - mortality).

**Figure S6:**
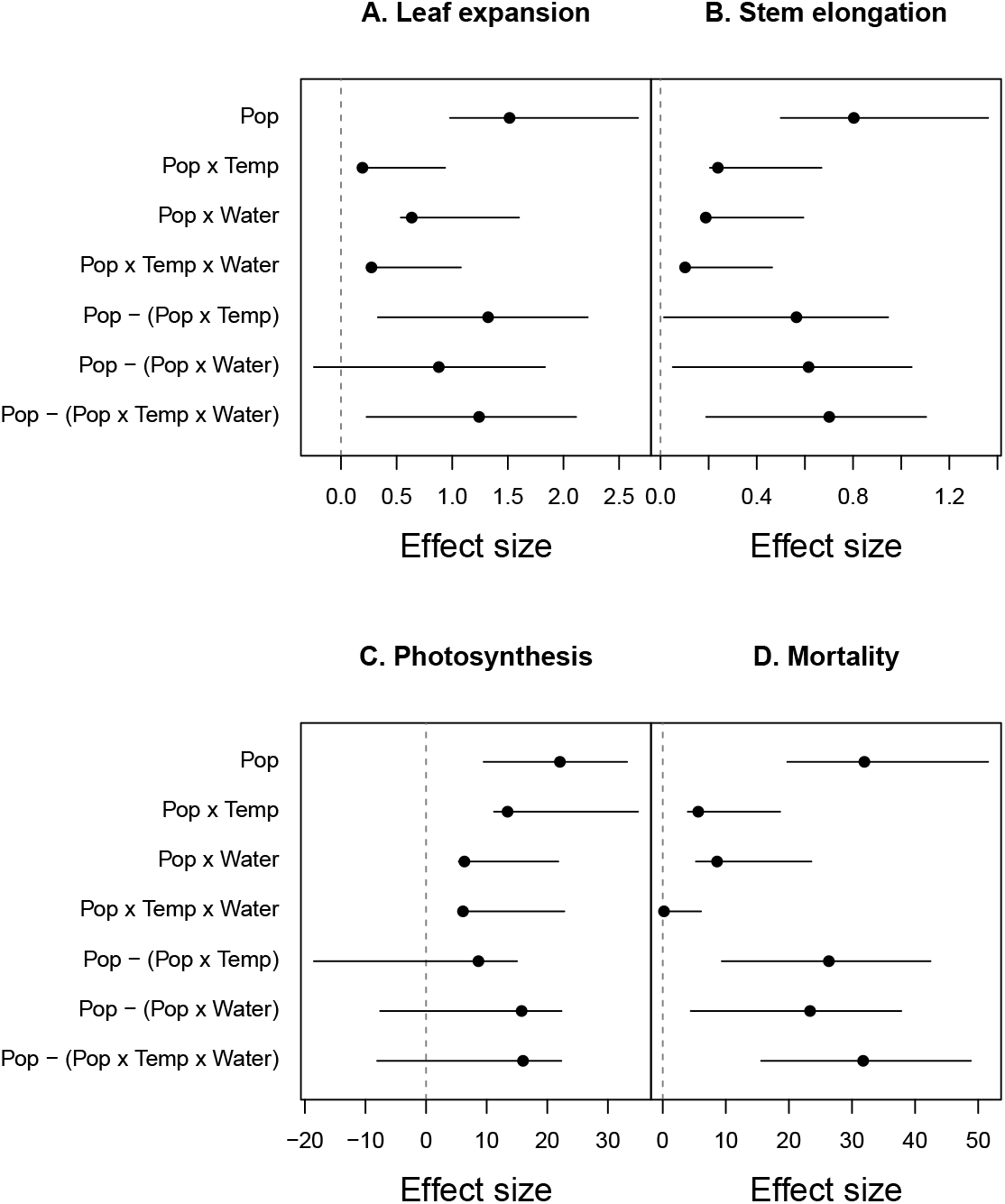
Population effect sizes are usually larger than Population × Treatment effect sizes. In each panel, we plot estimated effect size (points) and 95% confidence intervals (lines) inferred using parametric bootstrap (see Materials and Methods). At the top, we plot the effect sizes of Population (‘Pop’), two-way interactions between Population and Temperature (‘Pop × Temp’) or Water (‘Pop × Water’), and the three-way interaction between Population, Temperature, and Water (‘Pop × Temp × Water’). Below that we plot the difference in the Population minus the Population × Treatment effect size (e.g. ‘Pop - (Pop × Temp)’). When confidence intervals do no overlap zero (dashed line), this means that Population has a significantly greater effect size than the interaction. Effect sizes were measured using unstandarized mean square error for linear mixed-effects models (leaf expansion, stem elongation, and photosynthesis) and χ^2^ for GLMM (mortality). Hence, the effect size values are not comparable between different traits.

**Figure S7:**
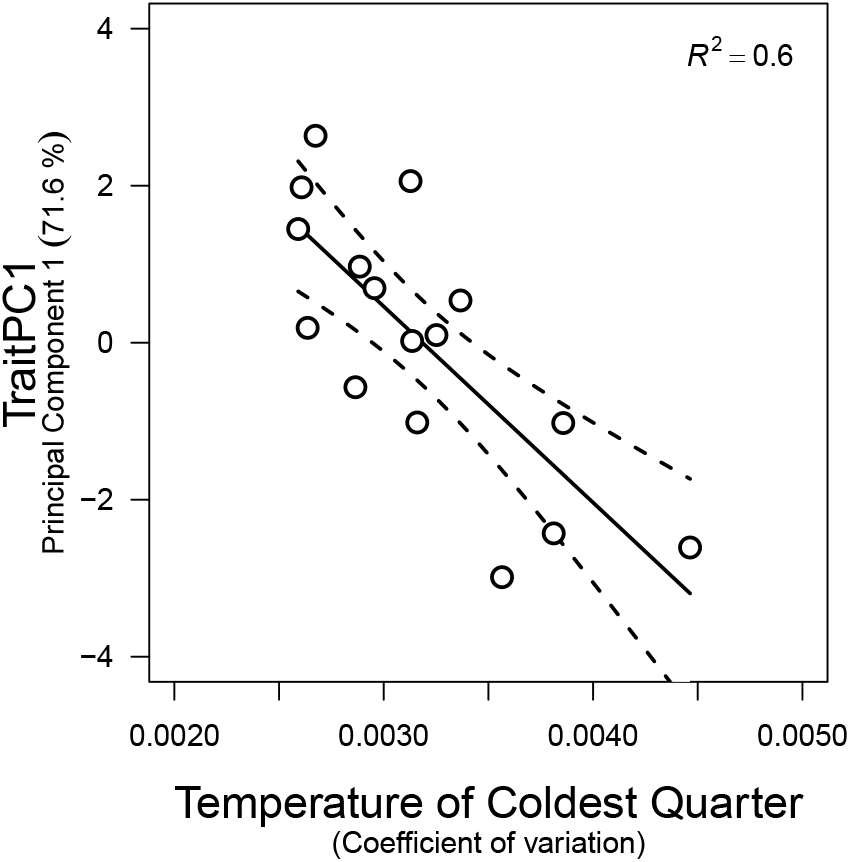
Trait variation, from fast to slow growth, is closely associated with neighborhood variation in temperature of the coldest quarter (bio11_σ_) Each point is a population coefficient of variation in bio11 averaged over a 62-km climatic neighborhood (x-axis) and position along the slow to fast growth axis (y-axis), defined as Principal Component 1 of four traits (see Material and Methods). The line and 95% confidence intervals were estimated using linear regression.

## Supporting Material and Methods

### Temperature treatments

We simulated typical growing season (June 1 - August 15) air temperatures at the two most thermally divergent focal sites in our study, Whitewater Canyon (WWC, Hot) and Little Jameson (LIJ, Cool). We downloaded daily interpolated mean, minimum, and maximum air temperature from 13 years (2000-2012) at both sites from ClimateWNA (Wang et al., 2012). This range was chosen because seeds used in the experiment were collected around 2012, thus their presence in that location at that time suggests that populations were able to persist there for at least some years before collection. Monthly temperatures from Cli-mateWNA are highly correlated with the air temperature recorded from data loggers in the field at these sites (A. Angert, unpub. data). Hence, the ClimateWNA temperature profiles are similar to actual thermal regimes experienced by *E. cardinalis* in nature. We simulated realistic temperature regimes by calculating the mean temperature trend from June to August using LOESS (Cleveland et al., 1992). The residuals were highly autocor-related at both sites (warmer than average days are typically followed by more warm days) and there was strong correlation (r = 0.65) between sites (warm days in WWC were also warm in LIJ). The ‘VARselect’ function in the vars package for R (Pfaff, 2008) indicated that a lag two Vector Autoregression (VAR(2)) model best captured the within-site autocorrelation as well as between-site correlation in residuals. We fit and simulated from the VAR(2) model using the package dse (Gilbert, 2014) in R. Simulated data closely resembled the autocorrelation and between-site correlation of the actual data. From simulated mean temperature, we next selected minimum and maximum daily temperatures. Mean, min, and max temperature were highly correlated at both sites. We chose min and max temperatures using site-specific fitted linear models between mean, max, and min temperature, with additional variation given by normally distributed random deviates with variance equal to the residual variance of the linear models. For each day, the nighttime (22:00 - 6:00) chamber temperature was set to the simulated minimum temperature. During the middle of the day, temperature was set to the simulated maximum temperature, with a variable period of transition between min and max so that the average temperature was equal the simulated mean temperature.

### Watering treatments

For watering treatments, we simulated two extreme types of streams where *E. cardinalis* grows. In the well-watered treatment, we simulated a large stream that never goes dry during the summer growing season. In the drought treatment, we simulated a small stream that has ample flow at the beginning of the season due to rain and snow melt, but gradually dries down through the summer. In both treatments, plants were bottom-watered using water chilled to 7.5°C. Plants in the well-watered treatment were fully saturated every two hours during the day. Watering in the drought treatment gradually declined from every two hours to every day between May 20 (36 days after sowing) and 10 June (57 days after sowing). Simultaneously, the amount of bottom-watering per flood decreased, such that only the bottom of the cone-tainers were wetted by the end of the experiment.

